# Significantly reduced, but balanced, rates of mitochondrial fission and fusion are sufficient to maintain the integrity of yeast mitochondrial DNA

**DOI:** 10.1101/2024.07.18.604121

**Authors:** Brett T. Wisniewski, Laura L. Lackner

## Abstract

Mitochondria exist as dynamic tubular networks and the morphology of these networks impacts organelle function and cell health. Mitochondrial morphology is maintained in part by the opposing activities of mitochondrial fission and fusion. Mitochondrial fission and fusion are also required to maintain mitochondrial DNA (mtDNA) integrity. In *Saccharomyces cerevisiae*, the simultaneous inhibition of mitochondrial fission and fusion results in increased mtDNA mutation and the consequent loss of respiratory competence. The mechanism by which fission and fusion maintain mtDNA integrity is not fully understood. Previous work demonstrates that mtDNA is spatially linked to mitochondrial fission sites. Here, we extend this finding using live-cell imaging to localize mtDNA to mitochondrial fusion sites. While mtDNA is present at sites of mitochondrial fission and fusion, mitochondrial fission and fusion rates are not altered in cells lacking mtDNA. Using alleles that alter mitochondrial fission and fusion rates, we find that mtDNA integrity can be maintained in cells with significantly reduced, but balanced, rates of fission and fusion. In addition, we find that increasing mtDNA copy number reduces the loss of respiratory competence in double mitochondrial fission-fusion mutants. Our findings add novel insights into the relationship between mitochondrial dynamics and mtDNA integrity.

## INTRODUCTION

In the majority of cell types, mitochondria exist as a dynamic network. The overall structure and distribution of the mitochondrial network are governed, in part, by the dynamic activities of mitochondrial fission and fusion. These activities are also critical to maintain mitochondrial function (Friedman and Nunnari, 2014; Lackner, 2014; Pernas and Scorrano, 2016; Chan, 2020; Giacomello *et al*., 2020). Mitochondrial fusion is thought to facilitate content mixing between mitochondrial compartments and is required for the maintenance of mitochondrial DNA (mtDNA). mtDNA encodes essential subunits of the respiratory chain and factors required for the expression of mtDNA genes. Mitochondrial fission plays critical roles in the distribution, transport, and quality control-mediated degradation of the organelle and mtDNA segregation. Together, the rates of mitochondrial fission and fusion determine the connectivity of the mitochondrial network (Sesaki and Jensen, 1999; Hoppins *et al*., 2007; Giacomello *et al*., 2020). Higher rates of mitochondrial fission relative to fusion result in less connected mitochondrial networks. In contrast, higher rates of mitochondrial fusion relative to fission result in more interconnected mitochondrial networks. The complete inhibition of mitochondrial fission or fusion drives mitochondrial morphology to the extremes of fragmentation or hyperfusion. The relative rates of mitochondrial fission and fusion can be altered in response to various biological contexts. For example, in mitosis in mammalian cells, fission-fusion dynamics favor fission and the fragmentation of the mitochondrial network is thought to facilitate the distribution and equal inheritance of mitochondria (Taguchi *et al*., 2007). In contrast, in stress conditions such as starvation, fission-fusion dynamics favor fusion and the hyperfused network promotes ATP production and stress resistance and protects mitochondria from autophagosomal degradation (Tondera *et al*., 2009; Gomes *et al*., 2011; Rambold *et al*., 2011).

When mitochondrial fission and fusion are simultaneously inhibited, mitochondrial networks have a near wild-type morphology but are functionally compromised (Sesaki and Jensen, 1999). In yeast, fission-fusion double mutants have defects in the maintenance of mtDNA integrity, are predicted to have a decreased rate of diffusion through the mitochondrial network, and exhibit increased sensitivity to stress and a reduced replicative lifespan (Rafelski *et al*., 2012; Bernhardt *et al*., 2015; Osman *et al*., 2015). In mice, defects in mitochondrial quality control are observed when fission and fusion are simultaneously disrupted (Song *et al*., 2017). Therefore, the effects of altered fission-fusion dynamics cannot be solely explained by alterations in the overall morphology of the mitochondrial network.

The core machineries driving mitochondrial fission and fusion are conserved from yeast to humans (Hoppins *et al*., 2007; Giacomello *et al*., 2020). The dynamin-related GTPase Dnm1/Drp1 (yeast/humans) is the core component of the mitochondrial fission machine. Dnm1/Drp1 assembles into helical structures that wrap around and mediate the scission of mitochondrial membranes (Ingerman *et al*., 2005; Mears *et al*., 2011). Dynamin-related proteins also serve as the major membrane remodelers for mitochondrial fusion. The DRPs Fzo1/Mitofusins and Mgm1/Opa1 are the primary drivers of outer and inner mitochondrial membrane fusion, respectively (Hoppins *et al*., 2007; Giacomello *et al*., 2020). Interestingly, the core fission and fusion machineries as well as the endoplasmic reticulum (ER) are present at both mitochondrial fission and fusion sites, suggesting coordinated regulation of the two processes (Murley *et al*., 2013; Lewis *et al*., 2016; Guo *et al*., 2018; Abrisch *et al*., 2020). The ER likely impacts the presence of factors, such as actin regulators, lipid-modifying enzymes, and/or elevated calcium concentrations, that help facilitate membrane remodeling at mitochondrial fission and fusion sites (Murata *et al*., 2020; Fung *et al*., 2023). In addition, ER-associated mitochondrial fission has been spatially linked to mtDNA in both yeast and mammalian cells and is proposed to couple mitochondrial fission to nucleoid distribution (Murley *et al*., 2013; Lewis *et al*., 2016). The spatial coupling of mtDNA to sites of mitochondrial fusion has yet to be demonstrated.

In budding yeast, the rates of mitochondrial fission and fusion are balanced making yeast an ideal model system to examine how alterations in fission-fusion dynamics impact mitochondrial function. Here, we use a series of established and novel fission-fusion mutants to determine the extent to which the mitochondrial fission-fusion balance can be perturbed while still maintaining mtDNA integrity.

## RESULTS AND DISCUSSION

### mtDNA is present at sites of mitochondrial fission and fusion but is not required to maintain mitochondrial fission and fusion rates

To further investigate the relationship between mitochondrial fission and fusion and mtDNA, we examined the spatial relationship between mtDNA and sites of mitochondrial fusion. To visualize mtDNA in live cells, we used the mt-LacO-LacI system developed by Osman et al., in which an array of LacO repeats are integrated into the mtDNA and bound by a mitochondria-targeted GFP-tagged LacI protein (Osman *et al*., 2015). We first confirmed the presence of mtDNA at sites of mitochondrial fission with the mt-LacO-LacI system in cells also expressing mitochondrial matrix-targeted dsRED (mito-RED). We observed mtDNA is associated with the majority (71%) of mitochondrial fission events (Figure 1, A and B). After mitochondrial fission, mtDNA was present in both of the newly generated tips in 68% of the events (Figure 1B). These results are consistent with those observed in Murley et al. (Murley *et al*., 2013). Similar to sites of mitochondrial fission, we found that mtDNA is associated with 77% of mitochondrial fusion events (Figure 1, A and C). Prior to the fusion event, mtDNA was present near the future fusion site in both fusion partners in the majority of events (73%) (Figure 1C).

**Figure 1.**
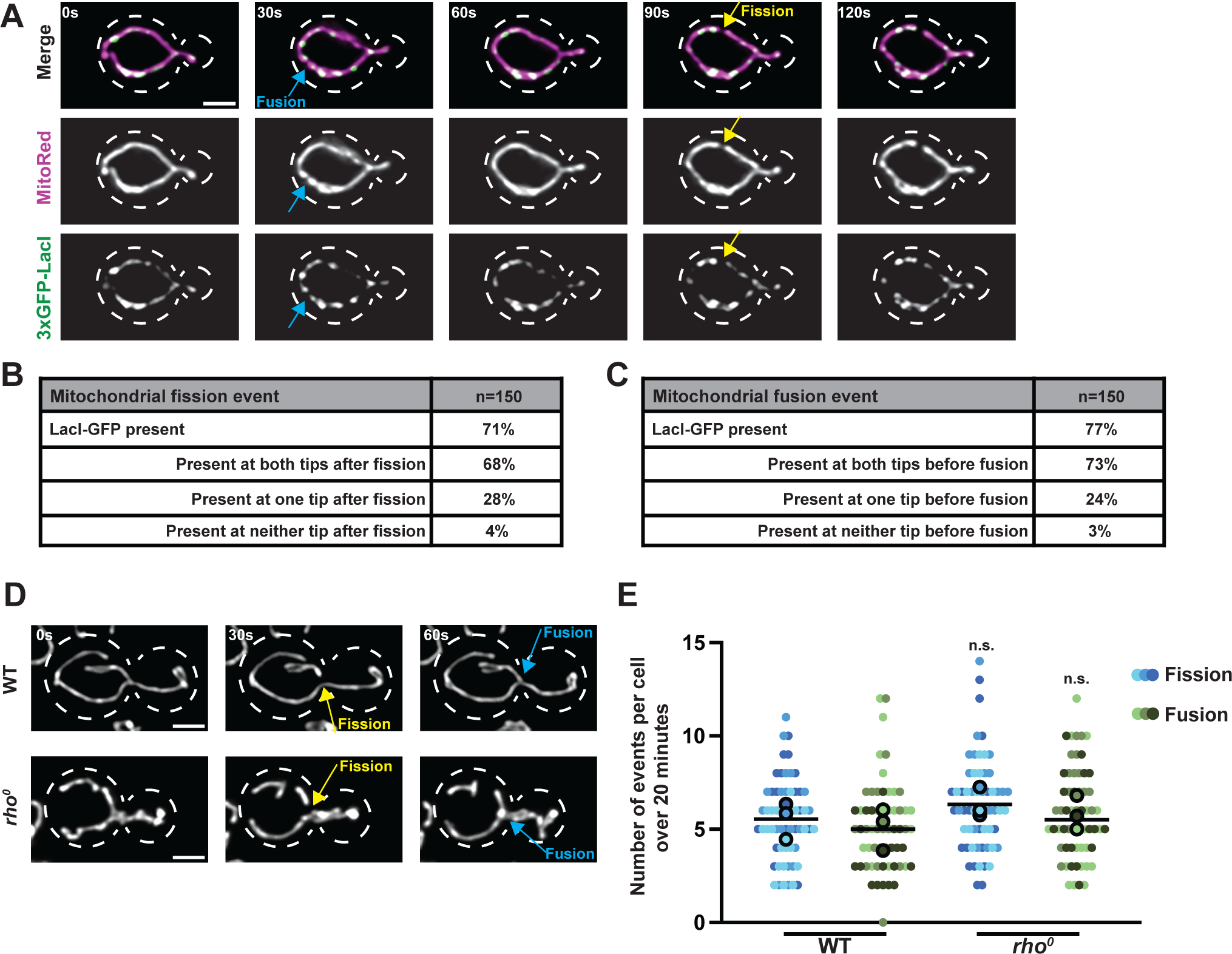
mtDNA is present at sites of mitochondrial fission and fusion but does not drive either process. (A-C) Wild-type cells expressing the mt-LacO-LacI system and mito-RED to visualize mtDNA and mitochondria, respectively, were grown in SCD media and analyzed by fluorescence microscopy. Yellow arrows indicate mitochondrial fission events and blue arrows indicate mitochondrial fusion events. Quantification of the presence of LacI-GFP in relation to fission and fusion sites is shown in panels C and D, respectively. (D) Wild-type and *rho^0^*cells expressing mito-RED were grown in SCD media and imaged by fluorescence microscopy. Yellow arrows indicate mitochondrial fission events and blue arrows indicate mitochondrial fusion events. (E) Quantification of mitochondrial fission and fusion events per cell in wild-type and *rho^0^* cells expressing mito-Red over twenty-minute time-lapse movies. Each dot represents a single cell. Twenty cells are quantified for each biological replicate, and the three biological replicates are represented in different colors. The black line denotes the grand mean. *p* values are in comparison to WT. n.s. not significant (unpaired t-test). For all images, maximum intensity projections are shown, and the cell cortex is outlined with a white dashed line. Bar, 2 µm.

We confirmed the presence of mtDNA at sites of mitochondrial fusion using a separate marker for mtDNA. GFP-tagged mtDNA binding proteins, such as Rim1 and Yme2, have been used to examine the behavior of mitochondrial nucleoids (mtDNA plus associated protein) (Murley *et al*., 2013; Friedman *et al*., 2015). To confirm the functionality of Rim1-GFP and Yme2-GFP, we examined the frequency with which cells expressing Rim1-GFP and Yme2-GFP lose respiratory competence using a standard petite frequency assay (see methods for details). We found that cells expressing Yme2-GFP had a petite frequency similar to that observed for wild-type cells, while the petite frequency of cells expressing Rim1-GFP was significantly increased (Figure S1A). Therefore, we moved forward using Yme2-GFP as a visual marker of mitochondrial nucleoids. We found Yme2-marked mitochondrial nucleoids were present at the majority of fission and fusion events (Figure S1, B-E). These results indicate that similar to mitochondrial fission, mtDNA is spatially associated with sites of mitochondrial fusion.

We next wanted to determine if the presence of mtDNA influenced the frequency of mitochondrial fission or fusion events. To this end, we quantified the number of fission and fusion events over twenty-minute time-lapse movies for both wild-type cells and *rho^0^* cells, which lack mtDNA. Consistent with Nunnari *et al*., we observed that the rates of mitochondrial fission and fusion were roughly balanced in wild-type cells (Figure 1, D and E) (Nunnari *et al*., 1997). Similar rates of mitochondrial fission and fusion were observed for *rho^0^*cells, indicating that the presence of mtDNA does not affect the frequency of mitochondrial fission and fusion events.

### Defects in mtDNA integrity are not specific to the loss of Dnm1 and Fzo1 but are a general consequence of simultaneous inhibition of fission and fusion

While the results above indicate that mtDNA is not required to maintain the mitochondrial fission-fusion balance, previous work suggests that the fission-fusion balance is required for the maintenance of mtDNA integrity (Osman *et al*., 2015). Specifically, cells lacking both Dnm1 and Fzo1 exhibit an increased frequency of becoming respiratory incompetent (i.e. petite). Interestingly, the respiratory incompetence of petite Δ*dnm1* Δ*fzo1* cells is due to mtDNA mutation (*rho^-^* cells) and not mtDNA loss (*rho^0^* cells) (Osman *et al*., 2015). To determine if the defect in mtDNA maintenance is specific to the loss of Dnm1 and Fzo1 or is a general consequence of simultaneously inhibiting fission and fusion, we examined mtDNA maintenance in different fission-fusion mutant combinations. Deletions of *FIS1* and *MGM1* were used as alternative approaches to inhibit fission and fusion, respectively (Mozdy *et al*., 2000; Tieu and Nunnari, 2000; Wong *et al*., 2000), and the petite frequency of individual mitochondrial fission gene deletions and fission-fusion double mutant combinations was determined. The petite frequency of individual fusion gene deletions could not be determined since fusion mutants lose mtDNA and cannot be propagated in media that requires respiration (Jones and Fangman, 1992; Guan *et al*., 1993; Hermann *et al*., 1998). We found that there is a significant increase in the percentage of cells that become respiratory incompetent (i.e. petite) for all fission-fusion double mutant combinations tested (Figure 2A). In addition, the increase for the fission-fusion double mutants (>51% petite) was far greater than that observed for the disruption of fission alone (∼10% petite) (Figure 2A) (Hanekamp *et al*., 2002; Cheng *et al*., 2008). Mitochondrial morphology across the fission-fusion double mutant combinations was similar, with the majority of cells displaying cortical wild-type networks and an increase in cells with more collapsed, simple networks (Figure 2, B-D). Also, the majority of the petite cells for all fission-fusion double mutant combinations were *rho^-^*, not *rho^0^*(Figure 2E), similar to what was previously observed for Δ*dnm1* Δ*fzo1* cells (Osman *et al*., 2015). These results indicate that the defect in the maintenance of mtDNA integrity that is observed for cells lacking Dnm1 and Fzo1 is not specific to the loss of Dnm1 and Fzo1 and is a general consequence of simultaneously inhibiting fission and fusion.

**Figure 2.**
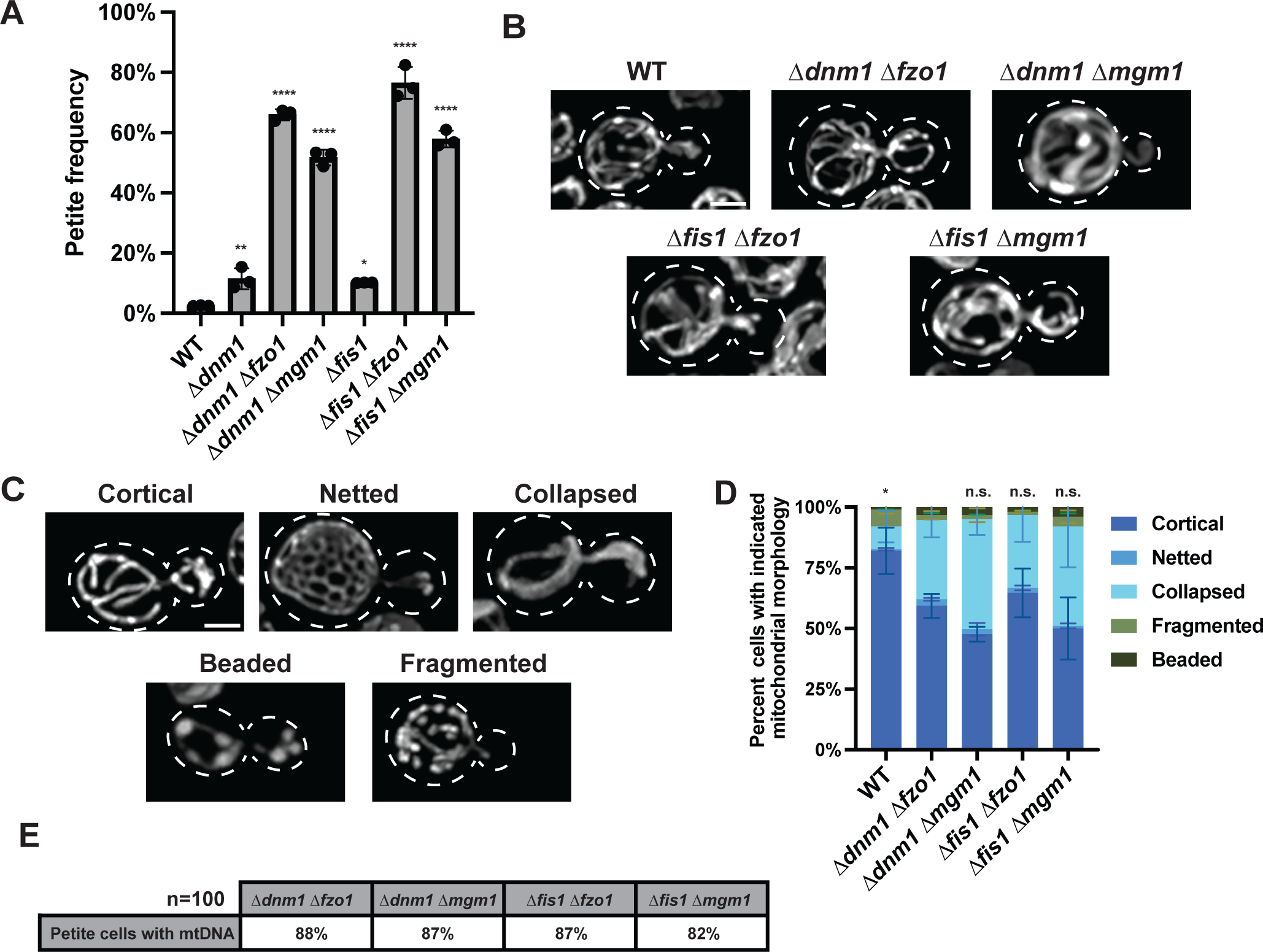
Defects in mtDNA integrity are a general consequence of simultaneous inhibition of mitochondrial fission and fusion. (A) The percentage of respiratory deficient cells determined by a petite frequency assay for the indicated genotypes. Three biological replicates are shown with each black dot representing the average for one biological replicate; >300 colonies were quantified per replicate. *p* values are in comparison to WT. * *p* < 0.1; ** *p* < 0.01; **** *p* < 0.0001; n.s. not significant (ordinary one-way ANOVA multiple comparisons). (B-D) Cells of the indicated genotype expressing mito-RED were grown in SCEG media and imaged by fluorescence microscopy. Panel B is a representative image of the cortical mitochondrial networks for each genotype. Panel C shows a representative cell for each mitochondrial morphology phenotype quantified in Panel D (n = three biological replicates, 100 cells quantified per replicate). *p* values are in comparison to Δ*dnm1* Δ*fzo1* cortical percentage. * *p* < 0.1; n.s. not significant (ordinary one-way ANOVA multiple comparisons). Maximum intensity projections are shown. The cell cortex is outlined with a white dashed line. Bar, 2 µm. (E) Quantification of the percentage of respiratory incompetent cells of the indicated genotype that contain mtDNA and were classified as *rho^-^* (see methods "assaying petite colonies for *rho^-^* vs. *rho^0^*" for details). Cells from 100 petite colonies were quantified for each genotype.

### Characterization of *DNM1* alleles that have reduced but balanced rates of mitochondrial fission and fusion

Our results in combination with the results of others indicate that, even though wild-type-like mitochondrial networks are maintained, there is an increase in the percentage of cells that become respiratory incompetent due to mtDNA integrity defects when both mitochondrial fission and fusion are inhibited (Figure 2) (Osman *et al*., 2015). These results raise the question of whether there is a minimal level of fission and fusion that is required to maintain the integrity of mitochondrial genomes. To address this question, we sought a way to decrease the rates of mitochondrial fission and fusion while maintaining a balance between the two processes.

We recently found that adding a GFP tag to the C-terminus of Dnm1 creates a hypomorphic *DNM1* allele. Mitochondria in cells expressing Dnm1-GFP from the endogenous *DNM1* locus have wild-type-like morphology; however, when Dnm1-GFP is combined with a mutant that attenuates mitochondrial fission, the fission defect is exacerbated (Harper *et al*., 2023). We were interested in more rigorously characterizing the hypomorphic *DNM1-GFP* allele as well as a *DNM1-GFP\** allele, which includes a C-terminal auxin-inducible degron (AID) tag in addition to GFP. To this end, we examined fission-fusion events over twenty-minute time-lapse movies in cells expressing Dnm1-GFP or Dnm1-GFP* along with mito-RED (Figure 3, A-C). As expected, both Dnm1-GFP and Dnm1-GFP* localized to sites of mitochondrial fission and fusion (Figure 3, B and C) (Abrisch *et al*., 2020). Interestingly, we found that the number of fission events was reduced in cells expressing Dnm1-GFP and Dnm1-GFP*, with the reduction in events being most severe for Dnm1-GFP* (Figure 3, A and A’; average fission events per cell over 20 minutes: WT=5.5, Δ*dnm1* Δ*fzo1*=0.2, *DNM1-GFP*=1.6, *DNM1-GFP*\*=0.3). A reduction in fission events was not surprising given the hypomorphic nature of the Dnm1-GFP allele (Harper *et al*., 2023). Surprisingly, however, the low mitochondrial fission rates were balanced with similarly low rates of fusion (Figure 3, A and A’; average fusion events per cell over 20 minutes: WT=5.1, Δ*dnm1* Δ*fzo1*=0.1, *DNM1-GFP*=1.3, *DNM1-GFP*\*=0.4). Thus, *DNM1-GFP* and *DNM1-GFP*\* represent novel alleles of *DNM1* that not only reduce mitochondrial fission but also reduce mitochondrial fusion, explaining why near wild-type mitochondrial morphology is maintained in Dnm1-GFP and Dnm1-GFP* cells.

**Figure 3.**
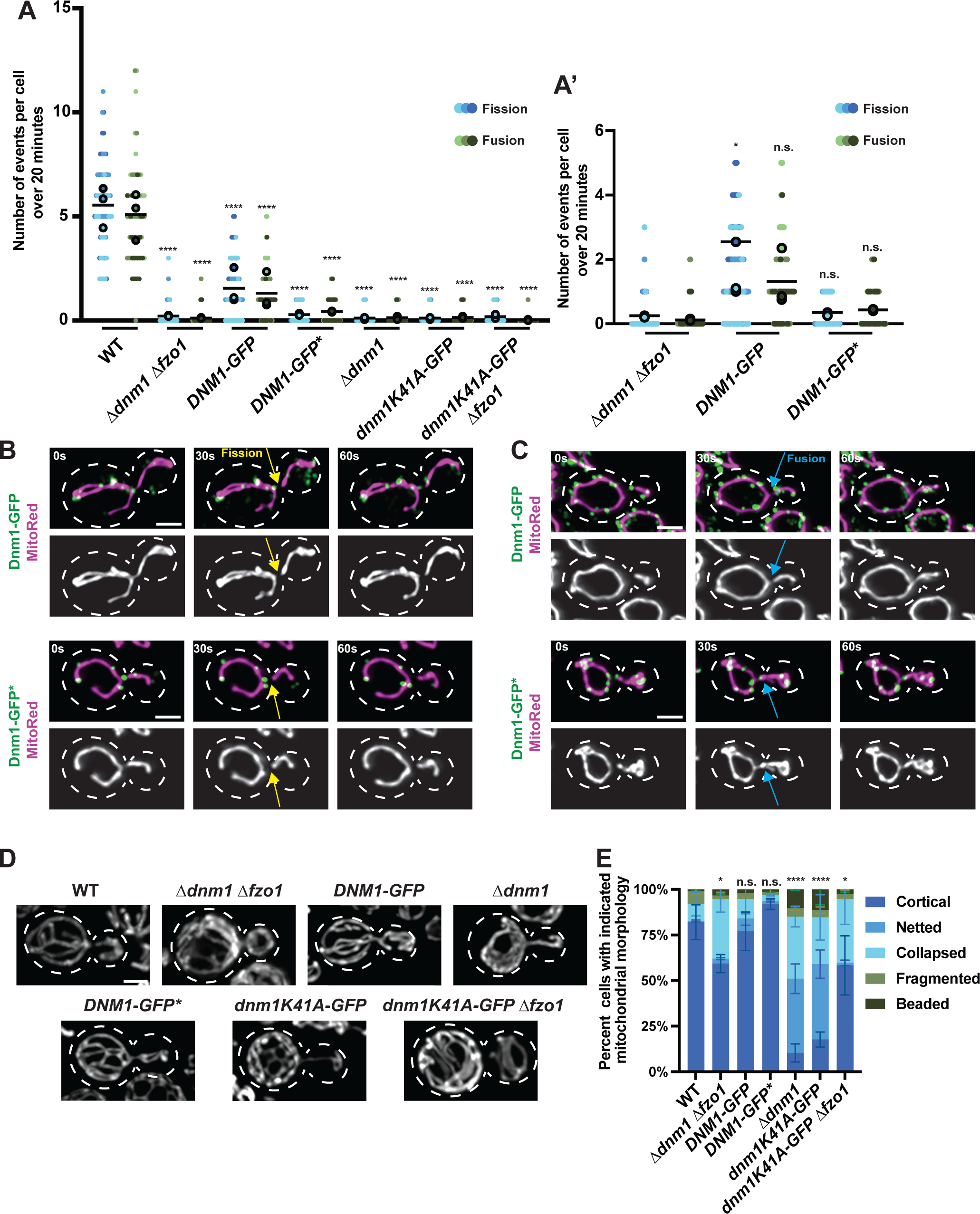
*DNM1* alleles reduce rates of mitochondrial fission and fusion while maintaining their balance. (A) Quantification of mitochondrial fission and fusion events per cell over a twenty-minute time-lapse movie of cells of the indicated genotype expressing mito-RED. Cells were grown in SCD media. Each dot represents a single cell. Twenty cells are quantified for each biological replicate, and the three biological replicates are represented in different colors. The black line denotes the grand mean. A’ is a magnified view of a subset of data shown in A. *p* values are in comparison to WT (A) and Δ*dnm1* Δ*fzo1* (A’). * *p* < 0.05; **** *p* < 0.0001; n.s. not significant (ordinary one-way ANOVA multiple comparisons). Data for WT are duplicated from 1E. (B and C) Cells expressing mito-RED and either Dnm1-GFP or Dnm1-GFP* were grown in SCD media and imaged by fluorescence microscopy. Yellow arrows indicate mitochondrial fission events in panel B. Blue arrows indicate mitochondrial fusion events in panel C. (D and E) Cells of the indicated genotype expressing mito-RED were grown in SCEG media and imaged by fluorescence microscopy. Panel D shows a representative image of the predominant mitochondrial morphology quantified for each genotype, with the full quantification of mitochondrial morphology for each genotype shown in Panel E (n = three biological replicates, 100 cells quantified per replicate). *p* values are in comparison to WT cortical percentage. * *p* < 0.05; **** *p* < 0.0001; n.s. not significant (ordinary one-way ANOVA multiple comparisons). Data for WT and Δ*dnm1* Δ*fzo1* are duplicated from 2D. For all images, maximum intensity projections are shown, and the cell cortex is outlined with a white dashed line. Bar, 2 µm.

These alleles differ from previously characterized Dnm1 mutants that have no fission activity, such as the GTP-hydrolysis deficient version of Dnm1, Dnm1^K41A^ (Naylor *et al*., 2006) (Figure 3, A-A’). In contrast to the predominantly wild-type-like cortical mitochondrial networks observed in cells expressing Dnm1-GFP and Dnm1-GFP*, cells expressing Dnm1^K41A^-GFP exhibit netted mitochondrial networks, similar to those seen in Δ*dnm1* cells (Naylor *et al*., 2006) (Figure 3, D and E). Thus, the hypomorphic *DNM1-GFP* and *DNM1-GFP\** alleles serve as novel and valuable tools to reduce mitochondrial fission and fusion while maintaining a balance in their rates.

### Significantly reduced, but balanced, rates of mitochondrial fission and fusion are sufficient to maintain mtDNA integrity

With the ability to maintain reduced, but balanced, rates of fission and fusion in *DNM1-GFP* and *DNM1-GFP*\* cells, we next asked whether mtDNA integrity could be maintained in these cells. We assessed mtDNA integrity by examining the percentage of cells that become respiratory incompetent using standard petite frequency assays. We found that in contrast to the high petite frequency of Δ*dnm1* Δ*fzo1* cells, *DNM1-GFP* and *DNM1-GFP\** cells have petite frequencies similar to wild-type cells (Figure 4A). The difference in respiratory competency between the strains was not a result of differences in mtDNA copy number; qPCR analysis revealed that the relative copy number of mtDNA was similar across the strains (Figure 4B). Together, these results indicate that significantly reduced, but balanced, rates of mitochondrial fission and fusion are sufficient to maintain mtDNA integrity.

**Figure 4.**
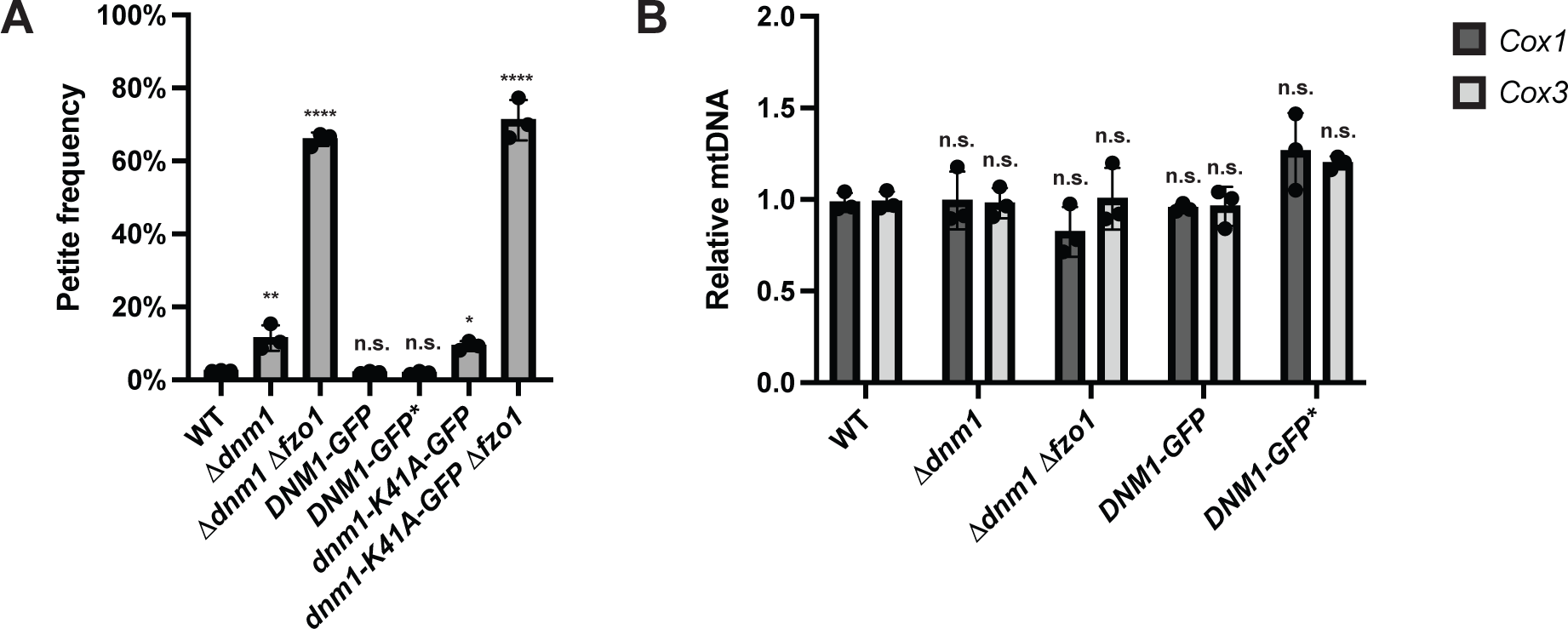
mtDNA can be maintained with significantly reduced, but balanced, rates of mitochondrial fission and fusion. (A) The percentage of respiratory deficient cells determined by a petite frequency assay for the indicated genotypes. Three biological replicates are shown with each black dot representing the average for one biological replicate; >300 colonies were quantified per replicate. *p* values are in comparison to WT. * *p* < 0.1; ** *p* < 0.01; **** *p* < 0.0001; n.s. not significant (ordinary one-way ANOVA multiple comparisons). Data for WT, Δ*dnm1*, and Δ*dnm1* Δ*fzo1* are duplicated from 2A. (B) Relative levels of mtDNA for cells of the indicated genotypes are shown. Quantitative PCR analysis was performed on DNA isolated from cells grown in YPEG. The level of mtDNA in cells for each genotype was normalized to the amount of mtDNA in wild-type cells. *p* values are in comparison to Δ*dnm1* Δ*fzo1*. n.s. not significant (ordinary one-way ANOVA multiple comparisons).

### Increased mtDNA copy number reduces the loss of respiratory competence in double mitochondrial fission-fusion mutants

We next sought to determine if there were ways to suppress the defect in cellular respiration in the absence of mitochondrial fission and fusion. Studies suggest that increased mtDNA copy number ameliorates defects associated with heteroplasmic mtDNA mutations (Nishiyama *et al*., 2010; Jiang *et al*., 2017; Filograna *et al*., 2019, 2021). Therefore, we wanted to test whether an increase in mtDNA copy number could rescue the respiratory competence of Δ*dnm1* Δ*fzo1* cells. Cells lacking the following proteins have been shown to have an increased mtDNA copy number (Figure S2A): Sml1, an inhibitor of the ribonucleotide reductase (Zhao *et al*., 1998; Taylor *et al*., 2005); Mrx6 and Mam33, proteins implicated in the maintenance of mtDNA copy number (Göke *et al*., 2020); and Cim1, a mitochondrial localized HMG-box protein that negatively regulates mtDNA copy number (Schrott and Osman, 2023).

We first determined that the loss of *SML1*, *MRX6*, *MAM33*, and *CIM1* in Δ*dnm1* Δ*fzo1* cells resulted in an increase in mtDNA copy number relative to Δ*dnm1* Δ*fzo1* cells (Figure 5A). We next examined the petite frequency of the triple mutant cells and observed that the loss of *SML1*, *MRX6*, *MAM33*, and *CIM1* in Δ*dnm1* Δ*fzo1* cells resulted in a decrease in the percentage of respiratory deficient cells (Figure 5B). Individual *SML1*, *MRX6*, *MAM33*, and *CIM1* deletions had petite frequencies similar to wild-type cells (Figure S2B). The rescue in petite frequency in the triple mutant cells was not due to alterations in mitochondrial morphology or dynamics; mitochondrial morphology and fission/fusion rates for the triple mutant cells and Δ*dnm1* Δ*fzo1* cells were similar (Figure 5, C-E). Together, these results indicate that the defects in respiratory competence resulting from defects in mtDNA integrity in Δ*dnm1* Δ*fzo1* cells can be partially suppressed by increasing mtDNA copy number, providing a possible model to understand how increased mtDNA copy number ameliorates mitochondrial dysfunction associated with heteroplasmic mtDNA mutations.

**Figure 5.**
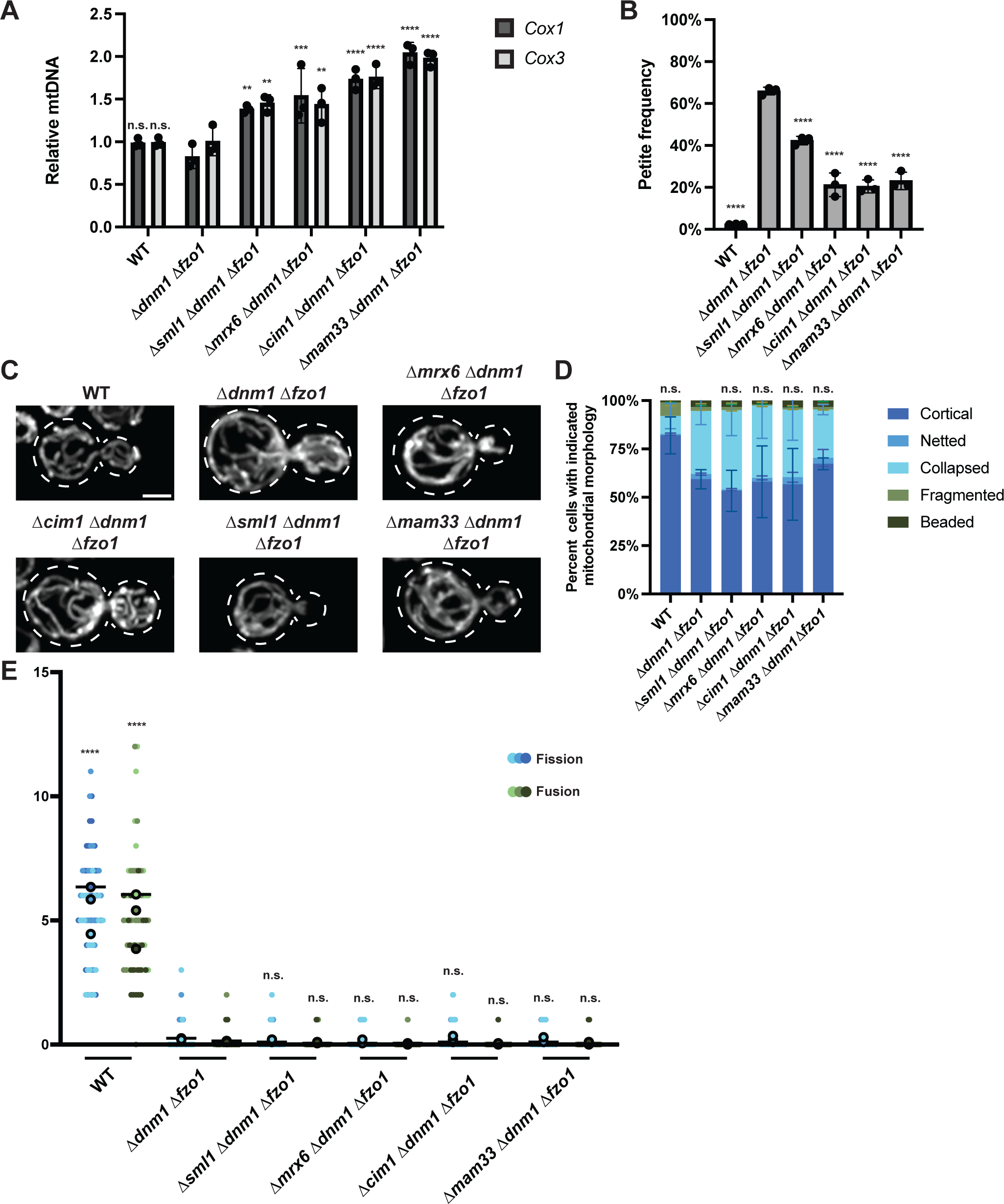
Increasing mtDNA copy number reduces the rate at which Δ*dnm1* Δ*fzo1* cells become respiratory incompetent. (A) Relative levels of mtDNA for cells of the indicated genotypes are shown. Quantitative PCR analysis was performed on DNA isolated from cells grown in YPEG. The level of mtDNA in cells for each genotype was normalized to the amount of mtDNA in wild-type cells. Three biological replicates are shown with each black dot representing the average for one biological replicate; 3 technical replicates were averaged for each biological replicate. *p* values are in comparison to Δ*dnm1* Δ*fzo1*. ** *p* < 0.01; *** *p* < 0.001; **** *p* < 0.0001; n.s. not significant (ordinary one-way ANOVA multiple comparisons). Data for WT and Δ*dnm1* Δ*fzo1* are duplicated from 4B. (B) The percentage of respiratory deficient cells determined by a petite frequency assay for the indicated genotypes. Three biological replicates are shown with each black dot representing the average for one biological replicate; >300 colonies were quantified per replicate. *p* values are in comparison to Δ*dnm1* Δ*fzo1*. **** *p* < 0.0001 (ordinary one-way ANOVA multiple comparisons). Data for WT and Δ*dnm1* Δ*fzo1* are duplicated from 2A. (C and D) Cells of the indicated genotype expressing mito-RED were grown in SCEG media and imaged by fluorescence microscopy. Panel C shows a representative image of the predominant mitochondrial morphology quantified for each genotype, with the full quantification of mitochondrial morphology for each genotype shown in Panel D (n = three biological replicates, 100 cells quantified per replicate). *p* values are in comparison to Δ*dnm1* Δ*fzo1* cortical percentage. n.s. not significant (ordinary one-way ANOVA multiple comparisons). Data for WT and Δ*dnm1* Δ*fzo1* are duplicated from 2D. Maximum intensity projections are shown. The cell cortex is outlined with a white dashed line. Bar, 2 µm. (E) Quantification of mitochondrial fission and fusion events per cell over twenty-minute time-lapse movies of cells of the indicated genotype expressing mito-RED. Cells were grown in SCD media. Each dot represents a single cell. Twenty cells are quantified for each biological replicate, and the three biological replicates are represented in different colors. The black line denotes the grand mean. *p* values are in comparison to Δ*dnm1* Δ*fzo1*. **** *p* < 0.0001; n.s. not significant (ordinary one-way ANOVA multiple comparisons). Data for WT and Δ*dnm1* Δ*fzo1* are duplicated from 3A.

In total, our findings add novel insights into the relationship between mitochondrial dynamics and mtDNA integrity. Our data indicate that despite mtDNA being spatially coupled to mitochondrial fission and fusion events, the rates of mitochondrial fission and fusion are not altered in cells lacking mtDNA. While mtDNA does not appear to affect mitochondrial dynamics, mitochondrial dynamics are required for the maintenance of mtDNA integrity for reasons not yet understood (Osman *et al*., 2015). Surprisingly, we found that reducing fission/fusion rates by ∼20-fold has no adverse effect on mtDNA integrity, suggesting that only a limited number of fission/fusion events are required for the factors/activities necessary for mtDNA maintenance. Additionally, we found the cellular respiration defect of fission-fusion double mutants could be partially suppressed by a modest (≤2-fold) increase in mtDNA copy number. Going forward, it will be critical to understand how significantly reduced, but balanced, rates of mitochondrial fission/fusion impact content mixing, molecular diffusion, and functional asymmetry within mitochondrial networks, which have been associated with quality control, cell fate determination, and aging (Chen and Chan, 2010; Giacomello *et al*., 2020; Klecker and Westermann, 2020).

## MATERIALS AND METHODS

### Strains and plasmids

Supplemental Tables 1 and 2 list all strains and primers used for this study, respectively. The wild type background strain was W303 (*ade2–1; leu2–3; his3–11, 15; trp1–1; ura3–1; can1–100*) (Thomas and Rothstein, 1989). All new yeast strains were generated via PCR-based targeted homologous recombination and transformation or mating, followed by sporulation and tetrad analysis. Primer sets used to amplify deletion cassettes are named F1 and R1 and primer sets used to amplify C-terminal gene tagging cassettes are named F5 and R3 and AID F and R (Longtine *et al*., 1998; Janke *et al*., 2004; Sheff and Thorn, 2004; Morawska and Ulrich, 2013). Plasmids used in this study are listed in Supplemental Table 3. For imaging mitochondria, strains were transformed with EcoRI digested pLL19 (pRS305 *mito-dsRed::LEU/NAT*)(Abrisch *et al*., 2020). To generate rho^0^ cells (Fox *et al*., 1991), cells were grown in yeast extract/peptone with 2% (wt/vol) dextrose (YPD) plus 25 µg/ml ethidium bromide for 24 h at 30°C and were used to inoculate a second culture in the same medium. Following another 24 h of growth, the culture was streaked on YPD plates. Single colonies were isolated and tested for lack of respiratory growth on yeast extract/peptone with 3% (vol/vol) ethanol and 3% (vol/vol) glycerol (YPEG) media. All strains and plasmids will be made available upon request.

### Petite frequency

The petite frequency assay takes advantage of the *ade2* mutation, which is present in our wild-type W303 strain background. *ade2* mutant cells that are respiratory competent turn red on media lacking adenine, while *ade2* mutant cells that lack functional mtDNA and, as a result, are respiratory incompetent, remain white (Shadel, 1999). Yeast cells were grown to log phase in yeast extract/peptone with 3% (vol/vol) ethanol and 3% (vol/vol) glycerol (YPEG) media at 30°C for 16-24 hours. Once in log phase, cells were diluted and spread on yeast extract/peptone with 2% (wt/vol) dextrose (YPD) plates not supplemented with adenine. Plates were incubated at 30°C for 2 days and stored at 4°C for one week for colony color to develop. Colonies were scored as red (*rho^+^*) or white (petite). For each genotype, >300 colonies were scored per biological replicate for a total of 3 biological replicates.

### Assaying petite colonies for *rho*^-^ vs. *rho*^0^

Cells from colonies that were scored as petite (see petite frequency) were lysed in 20 mM sodium hydroxide at 100°C for 10 minutes. Cell debris was pelleted by centrifugation and crude lysates were subjected to PCR analysis using primers specific to *ATP9*. Primers are listed in Supplemental Table 2. Colonies for which lysates produced a PCR product of the correct size were scored as *rho*^-^. Colonies for which lysates did not produce a PCR were subjected to DAPI analysis as a secondary screen to assess the presence of mtDNA (details can be found in the Imaging section). DAPI fluorescence was scored as either nuclear (indicating *rho*^0^) or mitochondrial (indicating *rho^-^*).

### Quantitative PCR

Real-time quantitative PCR (qPCR) was used to quantify the relative amounts of mtDNA in the indicated strains. Because the mtDNA of Δ*dnm1* Δ*fzo1* cells is prone to structural variations such as inversions and deletions in the vicinity of origins of replication (Osman *et al*., 2015), we chose to examine two different genes on the mtDNA, *COX1* and *COX3*, which are located in different regions of the mitochondrial chromosome. Total DNA was isolated from 5 mL of cultures grown to log phase in YPEG at 30°C using Biosearch Technologies MasterPure Yeast DNA Purification Kit for DNA extraction. DNA concentration of samples was determined by absorbance at 260 nm using a Molecular Devices SpectraMax iD3 multi-mode plate reader. Total DNA samples were diluted to 100 ng/mL. DNA was then subjected to qPCR analysis using Applied Biosystems PowerUp SYBR Green Master Mix and Applied Biosystems Quant studio 3 Real-Time PCR System. Primers specific for *COX1* and *COX3* were used for mtDNA and primers for *ACT1* were used for normalization to nuclear DNA. Primers are listed in Supplemental Table 2. Reactions were performed in Applied Biosystems MicroAmp Optical 96-Well Reaction Plates with Optical Adhesive Covers. Each reaction plate contained 2 biological WT controls for standardization. Each biological replicate consisted of 3 technical replicates, and a total of 3 biological replicates were quantified for each genotype. Relative mtDNA copy numbers were calculated using the comparative ΔΔCt method by determining Ct threshold values and using equations ΔΔCt = (Ct_COX2 − Ct_ACT1) at *t*(*n*) − (Ct_COX2 − Ct_ACT1) at *t*(0) and 2−ΔΔCt (Livak and Schmittgen, 2001).

### Imaging

Cells were grown to mid-log phase in either synthetic complete medium plus 2% (wt/vol) dextrose (SCD) with 2x adenine at pH 6.4 or synthetic complete medium plus 3% ethanol (vol/vol) and 3% glycerol (vol/vol) (SCEG) with 2x adenine at pH 6.4, as indicated. Cells were concentrated by centrifugation (3000 x g for 1 minute) and then imaged on a 4% (wt/vol) agarose pad on a depression slide. All cells were imaged on a Nikon Spinning Disk Confocal System fitted with a CSU-W1 dual-spinning disk unit (Yokogawa). A 60X (NA-1.42) objective and a Hamamatsu ORCA Fusion Digital CMOS camera were used for image capture. Images were captured and deconvolved with Elements software (Nikon). Linear adjustments to brightness and contrast were performed using FIJI (Schindelin *et al*., 2012). Deconvolved images are shown. Images of cells were cropped and assembled using Photoshop and Illustrator (Adobe System).

For mitochondrial morphology analysis (Figures 2B-D, 3D, 3E, 5C, 5D), all strains were grown in SCEG, and single time point images were captured. For mitochondrial fission and fusion analysis (Figures 1A-E, 3A-C, 5E, S1B, S1C-E), all strains were grown in SCD, and 20-min time-lapse movies with 30s intervals were captured.

For DAPI staining, cells were grown in SCD pH 6.4 media. DAPI was added to cultures to a final concentration of 2.5 µg/mL. Cells were incubated with DAPI for 30 minutes at room temperature, washed once with SCD, and then prepared and imaged as described above. DAPI fluorescence was scored as either nuclear (indicating *rho*^0^) or mitochondrial (indicating *rho^-^*).

### Image quantification

For mitochondria morphology quantification, single time point maximum projection images were analyzed on a per-cell basis; each cell’s mitochondrial network was classified as cortical, netted, collapsed, fragmented, or beaded. Examples of each classification are shown in Figure 2C. For each genotype, 3 biological replicates of 100 individual cells each were classified.

For mitochondrial fission and fusion event quantifications, twenty-minute time-lapse movies were analyzed and fission and fusion events were counted on a per-cell basis. For determining the presence of LacI-GFP or Yme2-GFP at sites of fission and fusion, 10 background fluorescence measurements from 5 x 5 pixel boxes were obtained for each movie quantified and used to set a threshold for signal intensity above background using maximum intensity value from among each 5 x 5 pixel box for the movie. Next, sites of fission or fusion were identified using the mitochondrial matrix marker signal only. 5 x 5 pixel boxes were then used to measure the intensity of GFP at these sites before and after each event as indicated in Figure 1. Instances in which the event site maximum intensity measurements were over background maximum intensity (determined as described above) were considered to have LacI-GFP or Yme2-GFP present. For each genotype, 3 biological replicates, each consisting of 100 fission and fusion events, were analyzed for the presence of LacI-GFP or Yme2-GFP at the sites of fission and fusion.

### Statistical analysis

Data distributions were assumed to be normal, and an unpaired t-test or a one-way ordinary ANOVA with Dunnett’s multiple comparisons was performed as indicated in the figure legend.

## ACKNOWLEDGEMENTS

We thank Jason Casler and members of the Lackner lab for feedback on the manuscript and critical scientific discussions. We also thank Northwestern’s Cell Biology Supergroup and the Wignall-Lackner Cell Biology Group for constructive feedback on the project. We thank Lauren Kraft and Gabriel Cavin-Meza for strain construction and Shelby Blythe and the Blythe lab for assistance with qPCR. Additional thanks to Jessica Hornick and Tong Zhang for assisting with the fluorescence microscopy. All microscopy was performed at the Biological Imaging Facility at Northwestern University (RRID:SCR_017767), supported by the Chemistry for Life Processes Institute, the Northwestern University Office for Research, the Department of Molecular Biosciences, and the Rice Foundation. B.T.W. was supported by the NIGMS Training Grant T32GM008061. L.L.L. is supported by NIGMS grant R01GM120303.

---

**Figure S1.**
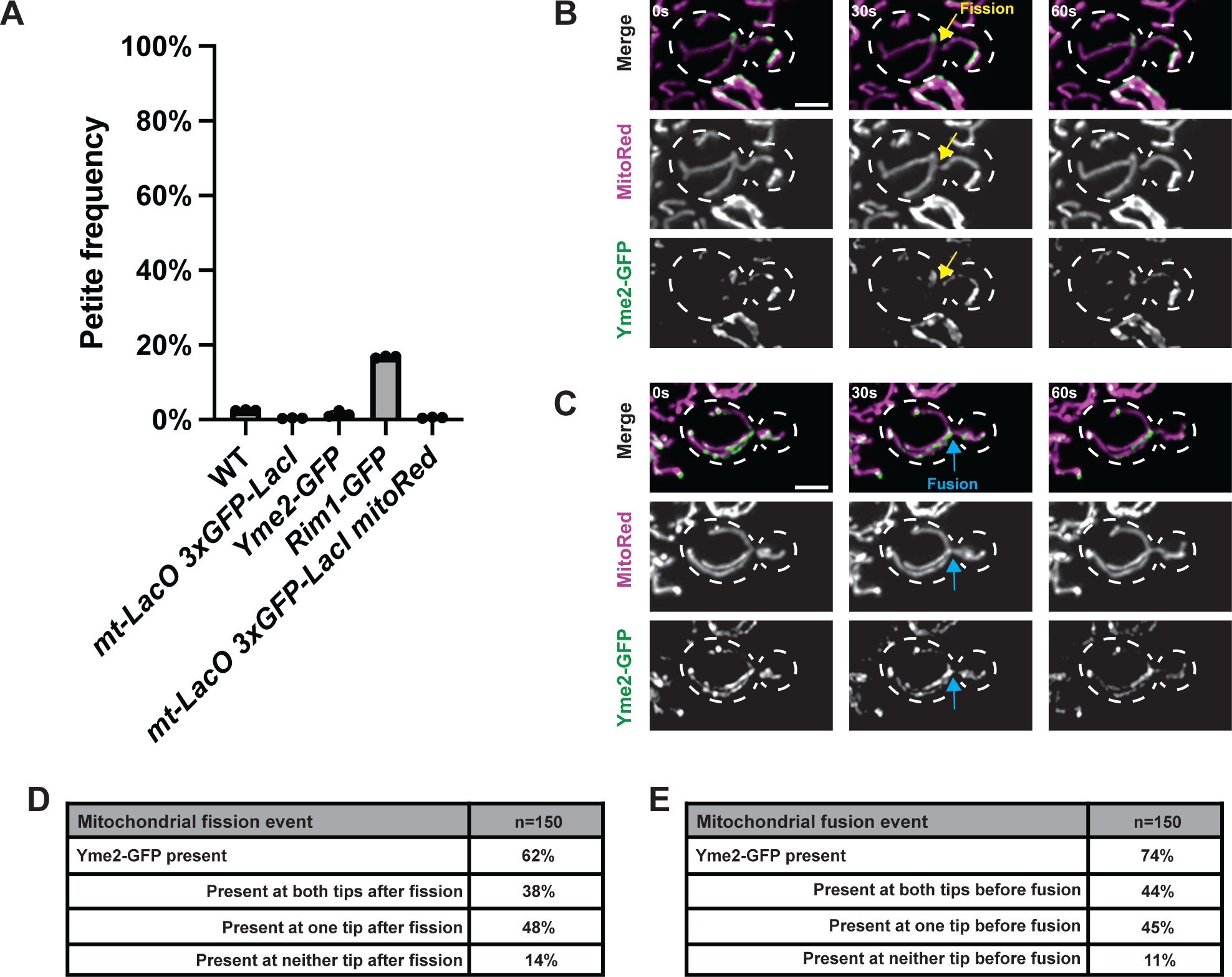
Mitochondrial nucleoids are present at sites of mitochondrial fission and fusion. (A) The percentage of respiratory deficient cells determined by a petite frequency assay for the indicated genotypes. Three biological replicates are shown with each black dot representing the average for one biological replicate; >300 colonies were quantified per replicate. Data for WT are duplicated from 2A. (B-E) Wild-type cells expressing the Yme2-GFP and mito-RED to visualize mitochondrial nucleoids and mitochondria, respectively, were grown in SCD media and analyzed by fluorescence microscopy. Maximum intensity projections are shown. The cell cortex is outlined with a white dashed line. Yellow arrows indicate mitochondrial fission events in panel B and blue arrows indicate mitochondrial fusion events in panel C. Bar, 2 µm. Quantification of the presence of Yme2-GFP in relation to fission and fusion sites is shown in panels D and E, respectively.

**Figure S2.**
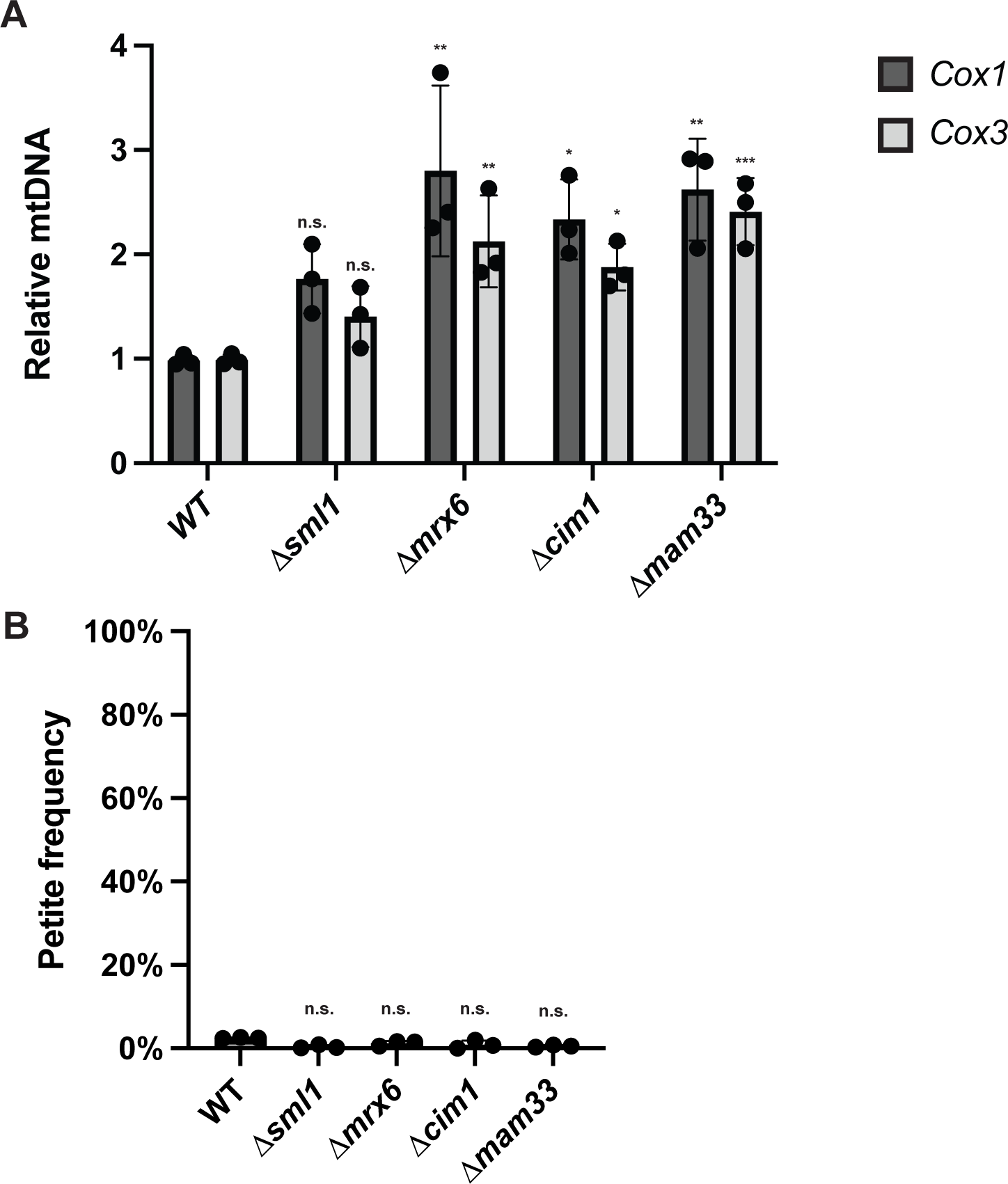
Relative mtDNA levels and petite frequency of Δ*sml1*, Δ*mrx6*, Δ*cim1*, and Δ*mam33* cells. (A) Relative levels of mtDNA for cells of the indicated genotypes are shown. Quantitative PCR analysis was performed on DNA isolated from cells grown in YPEG. The level of mtDNA in cells for each genotype was normalized to the amount of mtDNA in wild-type cells. p values are in comparison to WT. * *p* < 0.1; ** *p* < 0.01; *** *p* < 0.001; n.s. not significant (ordinary one-way ANOVA multiple comparisons). Data for WT are duplicated from 4B. (B) The percentage of respiratory deficient cells determined by a petite frequency assay for the indicated genotypes. Three biological replicates are shown with each black dot representing the average for one biological replicate; >300 colonies were quantified per replicate. p values are in comparison to WT. n.s. not significant (ordinary one-way ANOVA multiple comparisons). Data for WT and Δ*dnm1* are duplicated from 2A.

