## Supplemental Tables for "Significantly reduced, but balanced, rates of mitochondrial fission and fusion are sufficient to maintain the integrity of yeast mitochondrial DNA"

### Supplemental Material

Brett T. Wisniewski and Laura L. Lackner
Department of Molecular Biosciences, Northwestern University, Evanston, IL, 60208, USA.

B.T. Wisniewski: orcid.org/0000-0001-6157-1307
L.L. Lackner: orcid.org/0000-0003-0311-5199

**Supplemental Table 1: strains used in the study**

| LLY number | Genotype | Source |
|---|---|---|
| 92(A)/93(alpha) | W303 (*ade2–1; leu2–3; his3– 11,15; trp1–1; ura3–1; can1–100*) | Thomas and Rothstein, 1989 |
| 3220(alpha) | W303 *mt-LacO mt-3xGFP-LacI::KAN* pVT100-mito-dsRED | Osman et al., 2015 |
| 4025 (alpha) | W303 *mt-LacO mt-3xGFP-LacI::KAN* | This study |
| 4179 (alpha) | W303 *mt-LacO mt-3xGFP-LacI::KAN* | This study |
| 4734 | W303 *rho0 mitoRed:LEU/NAT* | This study |
| 3411(A)/3412(alpha) | W303 *mitoRed::LEU/NAT* | This study |
| 581 | W303 *Yme2-yEGFP::HIS* | This study |
| 666 | W303 *pRS304 Rim1-yEGFP* | This study |
| 3954 | W303 *Yme2-yEGFP::HIS mitoRed::LEU/NAT* | This study |
| 29(alpha)/30(A) | W303 *Δdnm1::NAT* | This study |
| 4451(A)/4452(alpha) | W303 *Δdnm1::NAT Δfzo1::NAT* | This study |
| 4031(A)/4032(alpha) | W303 *Δdnm1::kan Δmgm1::NAT* | This study |
| 4489/4490 | W303 *Δfis1::KAN* | This study |
| 4582(A)/4583(alpha) | W303 *Δfis1::KAN Δfzo1::NAT* | This study |
| 4584(A)/4585(alpha) | W303 *Δfis1::KAN Δmgm1::NAT* | This study |
| 4456 | W303 *Δfzo1* ::NAT *Δdnm1::KAN mitoRed:LEU/NAT* | This study |
| 4411(alpha)/4460(A) | W303 *Δdnm1::KAN Δmgm1::NAT mitoRed:LEU/NAT* | This study |
| 4647(A)/4648(alpha) | W303 *Δfis1::KAN Δfzo1::NAT mitoRed:LEU/NAT* | This study |
| 4412 | W303 *Δfis1::kan Δmgm1::NAT mitoRed:LEU/NAT* | This study |
| 3349(alpha)/1084(A) | W303 *Dnm1-yEGFP::HIS* | Harper et al., 2023 |
| 616 | W303 *Dnm1-AID-GFP::HYG* | This study |
| 2182 | W303 *Dnm1K41A-yEGFP::HIS* | This study |
| 4809/4810 | W303 *Dnm1K41A-yEGFP::HIS Δfzo1::NAT* | This study |
| 3348 | W303 *Dnm1-yEGFP::HIS mitoRed::LEU/NAT* | This study |
| 3244 | W303 *Δdnm1::NAT mitoRed::LEU/NAT* | This study |
| 3350 | W303 *Dnm1-AID-GFP::HYG mitoRed::LEU/NAT* | This study |
| 3243 | W303 *Dnm1K41A-yEGFP::HIS mitoRed::LEU/NAT* | This study |
| 6052/6053 | W303 *Dnm1K41A-yEGFP::HIS Δfzo1::NAT mitoRed::LEU/NAT* | This study |
| 5446(A)/5447(A) | W303 *Δsml1::HIS Δdnm1::NAT Δfzo1::NAT* | This study |
| 5116(A)/5117(alpha) | W303 *Δmrx6::HIS Δdnm1::NAT Δfzo1::NAT* | This study |
| 5444(A)/5445(alpha) | W303 *Δdpi34::HIS Δdnm1::NAT Δfzo1::NAT* | This study |
| 5916/5915 | W303 *Δmam33::HIS Δdnm1::NAT Δfzo1::NAT* | This study |
| 6069 | W303 *Δsml1::HIS Δdnm1::NAT Δfzo1::NAT mitoRed::LEU/NAT* | This study |
| 5997(A)/5998(alpha) | W303 *Δmrx6::HIS Δ*dnm1*::NAT Δfzo1::NAT mitoRed::LEU/NAT* | This study |
| 5999(A)/6000(alpha) | W303 *Δcim1::HIS Δdnm1::NAT Δfzo1::NAT mitoRed::LEU/NAT* | This study |

| | | |
|---|---|---|
| 6067/6068 | W303 Δ*mam33::HIS* Δ*dnm1::NAT* Δ*fzo1::NAT* *mitoRed::LEU/NAT* | This study |
| 5239(A)/5240(alpha) | W303 Δ*sml1::HIS* | This study |
| 5012(A)/5013(alpha) | W303 Δ*mrx6::HIS* | This study |
| 5237(A)/5238(alpha) | W303 Δ*cim1::HIS* | This study |
| 5843(A)/5844(A) | W303 Δ*mam33::HIS* | This study |
| 5286(A)/5287(alpha) | W303 Δ*dnm1::NAT* Δ*sml1::HIS* | This study |
| 5114(alpha)/5115(alpha) | W303 Δ*mrx6::HIS* Δ*dnm1::NAT* | This study |
| 5284(A)/5285(alpha) | W303 Δ*dnm1::NAT* Δ*cim1::HIS* | This study |
| 5922 | W303 Δ*mam33::HIS* Δ*dnm1::NAT* | This study |

**Supplemental Table 2: oligos used in the study**

| primer # | Sequence 5' to 3' | Purpose | |
|---|---|---|---|
| 310 | TGCCAGTAATGCAGTCATTACCAAATGTGAAGAAGAAATTAAAAACCTATCTAAGGGTGACGGTGCTGGTTTA | Tag *YME2* | F5 |
| 311 | GCAATATAGGGTTTACACATATGTACAGAAAAGCAGAATAGTAAGTTTTGTTTCATCGATGAATTCGAGCTCG | Tag *YME2* | R3 |
| 379 | GAATGGGAAGAAATTAGAAGATGCTGAGGGCCAAGAAAATGCTGCTTCTTCAGAAGGTGACGGTGCTGGTTTA | Tag RIM1 | F5 |
| 380 | CTCTAGATAAAAAATATCGAGGAAGAGTCGAAATAAGCAAGCGTAAATATTACAATCGATGAATTCGAGCTCG | Tag RIM1 | R3 |
| 125 | GAGTTTATCATTAAGTAGCTACCAGCGAATCTAAATACGACGGATAAAGACGGATCCCCGGGTTAATTAA | Delete *DNM1* | F1 |
| 126 | GCCCGCAATGTTGAAGTAAGATCAAAAATGAGATGAATTATGCAAGAATTCGAGCTCGTTTAAAC | Delete *DNM1* | R1 |
| 315 | CATAGCAACATTATCTGATATCACGGATAGAGGCAAAACGGTAGGCTCATTTAACGCGGATCCCCGGGTTAATTAA | Delete *FZO1* | F1 |
| 316 | GTAACATTATGTATATTGATTTGAAAAGACCTCATATATTTACAAGAATATGAATTCGAGCTCGTTTAAAC | Delete *FZO1* | R1 |
| 1522 | GGCCATCCCAAGAGTGGCGAACTATAACACATTAGTAAGGCGGATCCCCGGGTTAATTAA | Delete *MGM1* | F1 |
| 1523 | GCTATTTACAAATTCTCTAATGACACTATTTATTTTACAGAATTCGAGCTCGTTTAAAC | Delete *MGM1* | R1 |
| 133 | CGGCACATAGAAGCACAGATCAGAGCACAGCCATACAACATAAGTCGGATCCCCGGGTTAATTAA | Delete *FIS1* | F1 |
| 134 | CGATTCATTCTTATGTATGTACGTATGTGCTGATTTTTTATGTGCTTGGAATTCGAGCTCGTTTAAAC | Delete *FIS1* | R1 |
| 1243 | 5'-TTATGCAATTAGTATTAGCAGC-3' | Amplify *ATP9* | F |
| 1244 | 5'-CGAATAATAATAAGAATGAAACC-3' | Amplify *ATP9* | R |
| 463 | GAAATCACTCGGAGTTTATAAAAAGGCTGCAACCCTTATTAGTAATATTCTGGGTGACGGTGCTGGTTTA | Tag *DNM1* (GFP) | F5 |
| 464 | CTATAATCACGCCCGCAATGTTGAAGTAAGATCAAAAATGAGATGAATTATGCAATCGATGAATTCGAGCTCG | Tag *DNM1* (GFP) | R3 |
| 323 | AATCACTCGGAGTTTATAAAAAGGCTGCAACCCTTATTAGTAATATTCTGCGTACGCTGCAGGTCGAC | Tag *DNM1* (AID) | AID F |
| 324 | ATCACGCCCGCAATGTTGAAGTAAGATCAAAAATGAGATGAATTATGCAAATCGATGAATTCGAGCTCG | Tag *DNM1* (AID) | AID R |
| 793 | GAGTTTATCATTAAGTAGCTACCAGCGAATCTAAATACGACGGATAAAGAATGGCTAGTTTAGAAGATCTTATTCCTAC | Amplify *dnm1K41A* | F |
| 794 | CAGAATATTACTAATAAGGGTTGCAGCCTTTTTATAAACTCCGAGTGATTTC | Amplify *dnm1K41A* | R |

| | | | |
|---|---|---|---|
| 1855 | CTACAGATACAGCATTTCCAAGA | Amplify *COX1* for qPCR | F |
| 1856 | GTGCCTGAATAGATGATAATGGT | Amplify *COX1* for qPCR | R |
| 1235 | TTGAAGCTGTACAACCTACC | Amplify *COX3* for qPCR | F |
| 1236 | CCTGCGATTAAGGCATGATG | Amplify *COX3* for qPCR | R |
| 1237 | CACCCTGTTCTTTTGACTGA | Amplify *ACT1* for qPCR | F |
| 1238 | CGTAGAAGGCTGGAACGTTG | Amplify *ACT1* for qPCR | R |
| 1848 | CACTAACCTCTCTTCAACTGCTCAATAATTTCCCGCTCG GATCCCCGGGTTAATTAA | Delete *SML1* | F1 |
| 1849 | GGAAAGAGAAAAGAAAAGAGTATGAAAGGAACTGAATT CGAGCTCGTTTAAAC | Delete *SML1* | R1 |
| 1679 | GATCATTGCAGAAGTAGTGAGATTTAGTCGTACGTTAC GTCCGGATCCCCGGGTTAATTAA | Delete *MRX6* | F1 |
| 1680 | GAATATGTCACATAATAGAGCAGTGGATAAGTACTCAAT GCAAAAGAATTCGAGCTCGTTTAAAC | Delete *MRX6* | R1 |
| 1845 | CACTAACCTCTCTTCAACTGCTCAATAATTTCCCGCTCG GATCCCCGGGTTAATTAA | Delete *CIM1* | F1 |
| 1846 | GGAAAGAGAAAAGAAAAGAGTATGAAAGGAACTGAATT CGAGCTCGTTTAAAC | Delete *CIM1* | R1 |
| 1862 | GATACATAAACTATCACATATACTAACAAATAACAAATAC GGATCCCCGGGTTAATTAA | Delete *MAM33* | F1 |
| 1863 | CCCCGAGTCGCGAAATAAAGCAAAGGTATCTTGTTTCG AATTCGAGCTCGTTTAAAC | Delete *MAM33* | R1 |

**Supplemental Table 3: plasmids used in the study**

| LLEC # | Plasmid name | Source |
|---|---|---|
| 19 | pRS305 mitoRed | Abrisch et al., 2020 |
| 54 | pKT128 pFA6a–link–yEGFP–SpHIS5 | Sheff and Thorn, 2004 |
| 181 | pRS304 | Sikorski and Hieter, 1989 |
| 25 | pFA6-natMX3 | Goldstein and McCusker, 1999 |
| 27 | pFA6a-kanMX6 | Longtine et al., 1998 |
| 29 | pFA6a-His3MX6 | Longtine et al., 1998 |
| 413 | pRS304 *RIM1-GFP(S65T)* | Friedman et al., Elife 2015 |
| 404 | pHyg-AID*-GFP | Morawska and Ulrich, 2013 |
| 170 | pHS20 *dnm1K41A* | Naylor et al., 2006 |
